# The first archaeal PET-degrading enzyme belongs to the feruloyl-esterase family

**DOI:** 10.1101/2022.10.14.512230

**Authors:** Pablo Perez-Garcia, Jennifer Chow, Elisa Costanzi, Marno F. Gurschke, Jonas Dittrich, Robert F. Dierkes, Violetta Applegate, Golo Feuerriegel, Prince Tete, Dominik Danso, Julia Schumacher, Christopher Pfleger, Holger Gohlke, Sander H. J. Smits, Ruth A. Schmitz, Wolfgang R. Streit

## Abstract

Polyethylene terephthalate (PET) is a commodity polymer known to globally contaminate marine and terrestrial environments. Today, around 40 bacterial and fungal PET-active enzymes (PETases) are known, originating from four bacterial and two fungal phyla. In contrast, no archaeal enzyme has been identified to degrade PET. Here we report on the structural and biochemical characterization of PET46, an archaeal promiscuous feruloyl esterase exhibiting degradation activitiy on PET, bis-, and mono-(2-hydroxyethyl) terephthalate (BHET and MHET). The enzyme, found by a sequence-based metagenome search, was derived from a non-cultivated, deep-sea Candidatus Bathyarchaeota archaeon. Biochemical characterization demonstrated that PET46 is a promiscuous, heat-adapted hydrolase. Its crystal structure was solved at a resolution of 1.71 Å. It shares the core alpha/beta-hydrolase fold with bacterial PETases, but contains a unique lid common in feruloyl esterases, which is involved in substrate binding. Thus, our study significantly widens the currently known diversity of PET-hydrolyzing enzymes, by demonstrating PET depolymerization by a lignin-degrading esterase.

## INTRODUCTION

The global use of synthetic and fossil fuel-derived polymers on a multi-million-ton scale for over eight decades and the lack of concepts for recycling have led to an unprecedented accumulation of plastics of various sizes and blends in almost all ecological niches including the deep-ocean^1–5^. Plastic litter serves as a carrier for many microorganisms that can attach to their surface, constituting the so-called “plastisphere” ^6–8^. Many studies have described the microbial communities colonizing most commodity polymers such as polyethylene (PE), polypropylene (PP), or polystyrene (PS), but also polyethylene terephthalate (PET) or polyamides (PA), through 16S rDNA amplicon or metagenomic sequencing, and less often by FISH or DGGE analysis^8–11^. Most studies focused exclusively on bacterial lineages, while only a few identified eukaryotes or archaea in addition (*e.g.* Table 1 in ^12^). While it has been speculated that some of these attached microorganisms might potentially be involved in the degradation of the polymers, it is more likely that most of them will simply use the plastics as a biocarrier or metabolize the additives, but are not able to break down the polymers themselves^13,14^.

**Table 1:** PET46 has structural similarities to feruloyl esterases and bacterial PETases. Crystal structures included in the analysis in Figure 3 are sorted according to their Z-Score^69^ compared to PET46. FAE: Ferulic Acid Esterase/Feruloyl-Esterase. *Phylogeny could not be inferred.

| Protein Name | Protein Type | Host | Phylum | PDB | Z Score | RMSD | % id | Source |
| --- | --- | --- | --- | --- | --- | --- | --- | --- |
| PET46 | PETase FAE | Cand. Bathyarchaeota | Bathyarchaeota | 8B4U | 49.4 | 0.0 | 100 | This work |
| GthFAE | FAE | *G. thermoglucosidasius* | Firmicutes | 7WWH | 36.3 | 1.5 | 32 | <sup>76</sup> |
| Est1E | FAE | *B. proteoclasticus* | Firmicutes | 2WTM | 33.5 | 1.8 | 27 | <sup>38</sup> |
| LJ0536 | FAE | *L. johnsonii* | Firmicutes | 3PF8 | 32.8 | 1.9 | 28 | <sup>39</sup> |
| PET2 | PETase | N/A* | Proteobacteria* | 7ECB | 23.8 | 1.9 | 23 | <sup>23</sup> |
| Thc_Cut2 | PETase | *T. cellulosilytica* | Actinobacteria | 5LUJ | 23.5 | 1.7 | 22 | <sup>77</sup> |
| TfCut_2 | PETase | *T. fusca* | Actinobacteria | 4CG1 | 23.2 | 1.7 | 22 | <sup>78</sup> |
| Cut190 | PETase | *S. viridis* | Actinobacteria | 4WFK | 23.1 | 1.9 | 20 | <sup>79</sup> |
| Est119 | PETase | *T. alba* | Actinobacteria | 6AID | 23.0 | 1.7 | 22 | <sup>80</sup> |
| IsPETase | PETase | *I. sakaiensis* | Proteobacteria | 6EQE | 23.0 | 1.8 | 23 | <sup>81</sup> |
| PHL-7 | PETase | N/A* | Actinobacteria* | 7NEI | 22.9 | 1.9 | 21 | <sup>26</sup> |
| PLE629 | PETase | Marinobacter sp. | Proteobacteria | 7VPA | 22.9 | 1.8 | 23 | <sup>82</sup> |
| Thc_Cut1 | PETase | *T. cellulosilytica* | Actinobacteria | 5LUI | 22.9 | 1.7 | 22 | <sup>77</sup> |
| LCC | PETase | N/A* | Actinobacteria* | 4EB0 | 22.6 | 1.8 | 23 | 20 |
| RgPETase | PETase | *R. gumimpiphilus* | Proteobacteria | 7DZT | 22.4 | 1.8 | 23 | 83 |
| PET30 | PETase | *K. jeonii* | Bacteroidetes | 7PZJ | 21.9 | 1.9 | 23 | 27 |
| PE-H | PETase | *P. aestusnigri* | Proteobacteria | 6SBN | 21.5 | 2.1 | 25 | 22 |
| PLE628 | PETase | *Marinobacter* sp. | Proteobacteria | 7VMD | 20.2 | 2.1 | 26 | 82 |
| PmC | PETase | *P. mendocina* | Proteobacteria | 2FX5 | 19.9 | 2.2 | 19 | 84 |
| CalB | Lipase | *C. albicans* | Ascomycota | 1TCA | 16.6 | 2.9 | 12 | 28 |
| IsMHETase | MHETase | *I. sakaiensis* | Proteobacteria | 6QZ4 | 14.6 | 3.2 | 13 | 37 |
| HiC | PETase | *T. insolens* | Ascomycota | 4OYL | 12.9 | 3.1 | 12 | 28 |
| FsC | PETase | *F. solani* | Ascomycota | 1AGY | 12.7 | 3.2 | 11 | 28 |

Nevertheless, in recent years, several studies have identified microbial enzymes that are able to degrade some of these synthetic polymers, including PET, polyurethane (PUR), PA, and a few others from mainly renewable sources^13,15^. To date, approximately 120 enzymes have been described to act on these polymers (PAZy database^16^), most of them being esterases, amidases, and oxygenases. Many of these proteins have relatively low conversion rates, show promiscuous activity or are only active on oligomers. Even though some euryarchaea (*e.g.* Thermoplasmatales) and TACK-archaea (*e.g.* Thaumarchaeota, Crenarchaeota) have been found to colonize plastic particles of various sizes^17,18^, not a single plastic-active enzyme of archaeal origin has yet been identified to break down a synthetic polymer.

In the case of PET, the vast majority of degrading enzymes derive from bacteria, including Actinomycetota/Actinobacteria^19–21^, Pseudomonadota/Proteobacteria^22–24^, Bacillota/Firmicutes^25,26^ and recently Bacteroidota/Bacteroidetes^27^. Few enzymes have been identified in fungi (Eukarya), including *Candida antarctica* CalB, *Humicola insolens* HiC, and *Fusarium solani* FsC^28^. These enzymes share some common features: They are cutinases or esterases, their catalytic triad comprises Ser-Asp-His, the active site is fairly exposed to the solvent, and they are deprived of any lid domain. Furthermore, aromatic (Trp, Phe, Tyr) and Met residues within the catalytic pocket contribute to the binding of PET and the formation of the oxyanion hole^13,29,30^.

Within this work, we describe and characterize PET46 (NCBI accession RLI42440.1), the first enzyme from archaeal origin reported to hydrolyze PET polymer. The enzyme is encoded in the metagenome-assembled genome (MAG) of the Candidatus Bathyarchaeota archaeon B1_G2, a member of the TACK group that was found at the Guaymas Basin^31^. The experimentally established crystal structure of the protein is similar to bacterial PETdegrading enzymes, but reveals several unique features. These include differences in the amino acid composition in and around the active site compared to its bacterial and eukaryotic counterparts. Furthermore, the enzyme’s structure shows high homology to feruloyl esterases and a flexible lid domain covers its active site, which was not previously described elsewhere for PETases. These findings demonstrate higher diversity of PET-active enzymes and strongly suggest that more enzymes could be able to degrade PET, which possibly have not yet been discovered by strict sequence and structural homology searches.

## RESULTS

### Profile Hidden Markov Model (HMM) search identifies the novel archaeal PETase PET46

Previous research identified PET esterases in bacteria and two fungal phyla (Figure 1). Since hitherto no archaeal PETase had been described, we speculated that archaeal esterases might as well be capable to hydrolyze PET. To address this question, we performed global database searches using publicly available microbial genomes and metagenomes from NCBI’s non-redundant database using a previously published HMM-based search approach^23,32^.

**Figure 1:**
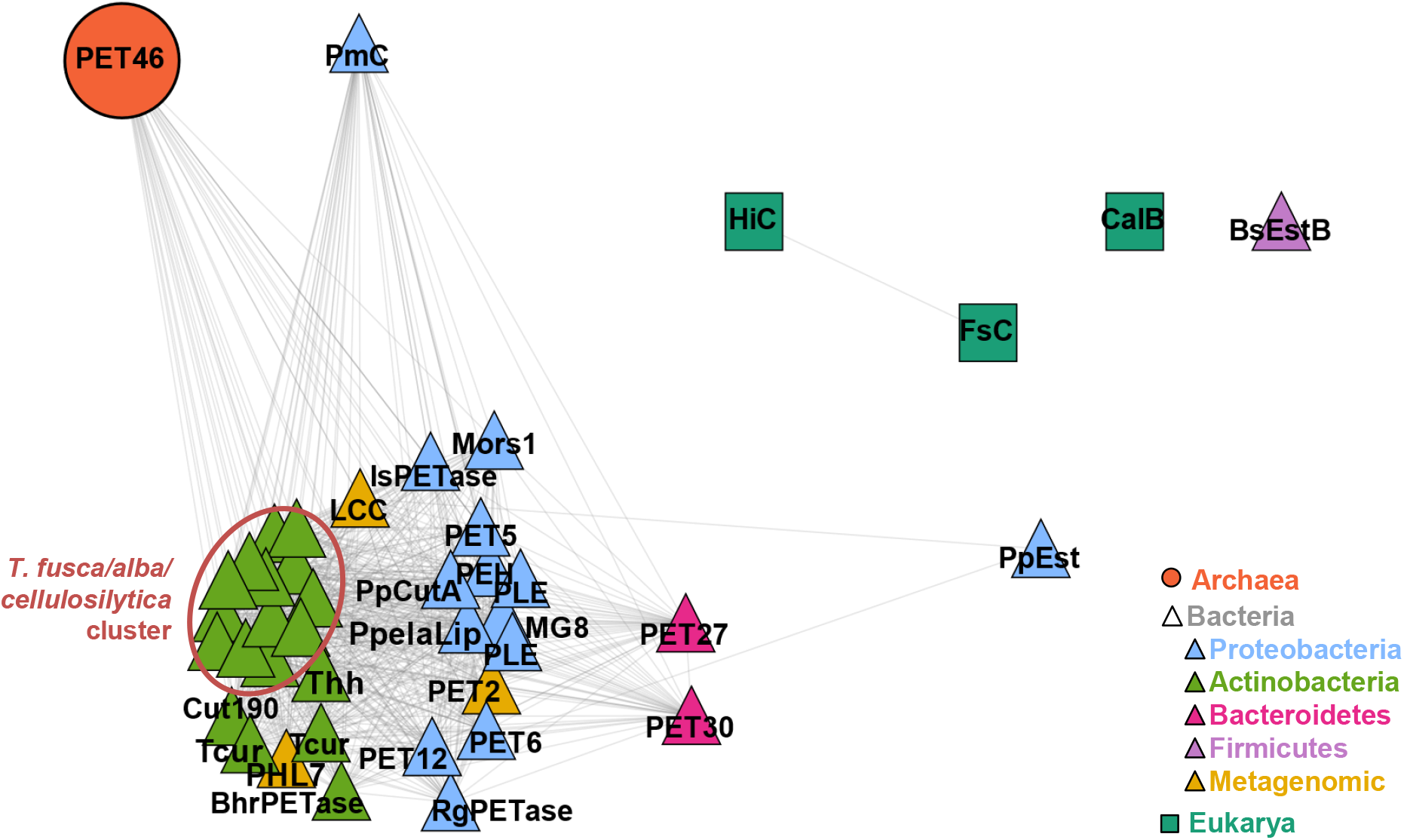
The “third domain” of PET degradation. The amino acid sequence of the first archaeal PET-degrading enzyme PET46 (coral orange, circle) was included in a sequence network analysis with all other known PETases from Bacteria (triangles) and Eukarya (squares) collected in PAZy^16^. The edge length between two nodes is inversely proportional to the BLASTp bitscore of both nodes (e-value < 0.05).

We selected PET46 as a putative archaeal PET-degrading esterase (Figure 1). Its sequence originates from a recently identified TACK archaeon found at a deep-sea marine sediment from the Guaymas Basin (Gulf of California, Mexico). The host strain Candidatus Bathyarchaeota archaeon B1_G2 has not been cultivated, but is part of a genome reconstruction project^31,33^. PET46 is encoded on a short 3,316 bp contig (QMYN01000264.1) by a 786 bp ORF coding for an alpha/beta-hydrolase (RLI42440.1) with 262 aa and a predicted molecular weight of 29.4 kDa (Supplementary Fig. S1). The other ORFs in the contig code for a tRNA-deacylase and two ribosomal proteins (Supplementary Fig. S1).

### Amino acid sequence and structural analyses imply that PET46 is a feruloyl esterase

For an initial characterization, the PET46 amino acid sequence was subjected to a more detailed bioinformatics analysis. Thereby, we identified a predicted G-x-S-x-G motif which is a common trait of serine hydrolases^34^. Amongst others, we identified conserved domains of dipeptidyl aminopeptidase/acylaminoacyl peptidase (DAP2), acetyl xylan esterase (AXE1), dienelactone hydrolase (DLH) and lysophospholipase (PldB, Supplementary Fig. S1). A BLASTp search against the non-redundant database results in 105 hits (query cov. > 80%, seq. id > 40%), from which only 26 are archaeal homologs, exclusively from the TACK-group. Most of them derive from Bathyarchaeota, and only three hits are related to the phylum Thermoproteota/Crenarchaeota. Interestingly, 79 homologs were found in Bacteria, especially in Firmicutes (Supplementary Fig. S1).

For further characterization, we produced PET46 wildtype (WT) heterologously in *E. coli* BL21 (DE3) carrying an N-terminal 6xHis-tag. The recombinant and purified protein was used for crystallization and additional biochemical tests.

Crystals of PET46 were obtained by sitting-drop vapor diffusion after 3-4 weeks. They were harvested, cryoprotected, flash-frozen in liquid nitrogen, and datasets were collected at the ESRF beamline ID23-1 (Grenoble, France). The best PET46 crystal grew in space group P 6’ 2 2 and diffracted to a resolution of 1.71 Å (Supplementary Table 1). We could unambiguously model the protein chains in the electronic density between residues 1-269. The final model was refined to Rwork/Rfree values of 15.23/17.27, and deposited to the PDB with accession ID 8B4U. All data collection and refinement statistics are reported in Supplementary Table 1.

One monomer is present in the asymmetric unit (ASU), which shares the common fold of the alpha/beta hydrolase superfamily, with the core of the enzyme being composed by eight β-strands connected by seven α-helixes (Supplementary Fig. S2). In addition, a lid domain composed by three α-helixes and two anti-parallel β-strands is present (Leu141-Val186). The active site is composed of the catalytic triad Asp206, His238, and Ser115. Interestingly, an unexpected electron density was present near the active site and it was modelled with a phosphate ion and two ethylene glycol molecules (Supplementary Fig. S2), likely coming from the protein buffer, crystallization solution and cryoprotectant.

Despite the low sequence similarity of only 23%, the structure of PET46 overlays the IsPETase from *Ideonella sakaiensis* (PDB 6EQE) with 1.8 Å Cα-RMSD (Figure 2 and Table 1). The largest difference is the medium-sized lid comprising 45 aa in PET46 (Leu141-Val186, Figure 2). Further structural differences around the active site are found in the enlarged loop between β4 and α3 (Loop 1; Asp68-Glu78; deep blue in Figure 2), which folds back to the outside, and the shorter loop between β10 and α10 containing the catalytic His (Loop 2; Arg234-Arg242; magenta in Figure 2). Loop 2 in IsPETase also contains one Cys that forms a disulfide bridge, which PET46 lacks. Almost all residues needed to form the oxyanion hole and the aromatic clamp are conserved or have similar properties as in other PETases^13^. Nevertheless, the lack of an equivalent to Trp185 in IsPETase suggests that the lid domain is involved in substrate binding and formation of the aromatic clamp (Figure 2). To answer this question, we constructed a chimera named PET46Δlid, where we substituted Ala140-Pro187 with the homologous Trp185-Thr189 minimal loop of the IsPETase. By this, we included the Trp185 involved in the formation of the aromatic clamp, which is missing in PET46 (Figure 2).

**Figure 2:**
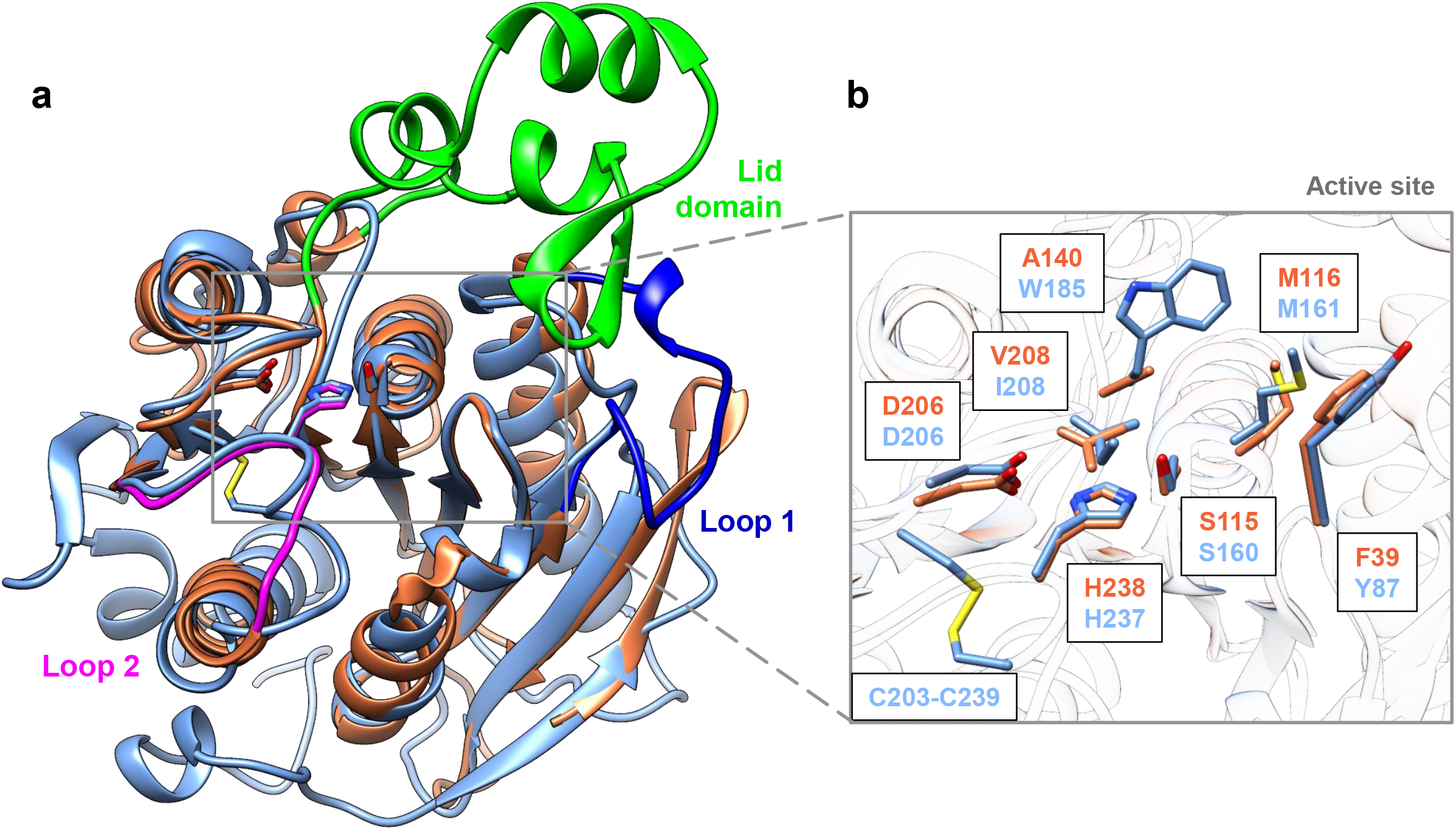
The crystal structure of PET46 resembles the crystal structure of the IsPETase - with unique features. Both proteins present the α/β-hydrolase fold and the same catalytic triad, but PET46 (coral orange; PDB 8B4U) presents a lid domain (bright green) that is not present in the IsPETase (sky blue; PDB 6EQE). Other structural differences are present in Loop 1 (deep blue) and Loop 2 (magenta) containing the active site His (**a**). The bacterial and the archaeal enzymes present the typical residues of Ser-hydrolases at the catalytically active positions (Ser, His and Asp), but PET46 lacks a Trp associated with PET binding and formation of the aromatic clamp in the IsPETase. Furthermore, PET46 also lacks a disulfide bridge in Loop 2 (**b**).

We further compared the structure of PET46 to all published bacterial and eukaryotic PETases (Table 1). Additionally, we performed searches against all crystal structures in the PDB. From this database, the best hits obtained were the feruloyl esterases GthFAE from *Geobacillus thermoglucosidasius* (PDB 7WWH) and Est1E from the rumen bacterium *Butyrivibrio proteoclasticus* (PDB 2WTM) together with the cinnamoyl esterase LJ0536 from *Lactobacillus johnsonii* (PDB 3PF8; Figure 3 and Table 1). All hits derive from Firmicutes. Feruloyl esterases are also known as ferulic acid esterases (FAEs). They are involved in plant biomass degradation and cleave *e.g.* cinnamic, *p*-coumaric or ferulic acid, thus “decoupling” plant cell wall polysaccharides and lignin^35^. Using ethyl cinnamate (EC) as a model substrate, we could detect enzyme-mediated H^+^ release derived from ester hydrolysis (Supplementary Fig. S3). These aromatic acids esterified to long polymers may be analogous to MHET units at the end of a PET chain (Figure 3 and Supplementary Fig. S3). FAEs are believed to be secreted enzymes, even if no apparent N-terminal signal peptide is present^36^. In the case of PET46, no obvious secretion signal is detected. Since FAEs form a protein family with tannases, to which the MHETase from *I. sakaiensis* belongs^37^, we also included its structure (PDB 6QZ4, Table 1) in our structural analysis.

**Figure 3:**
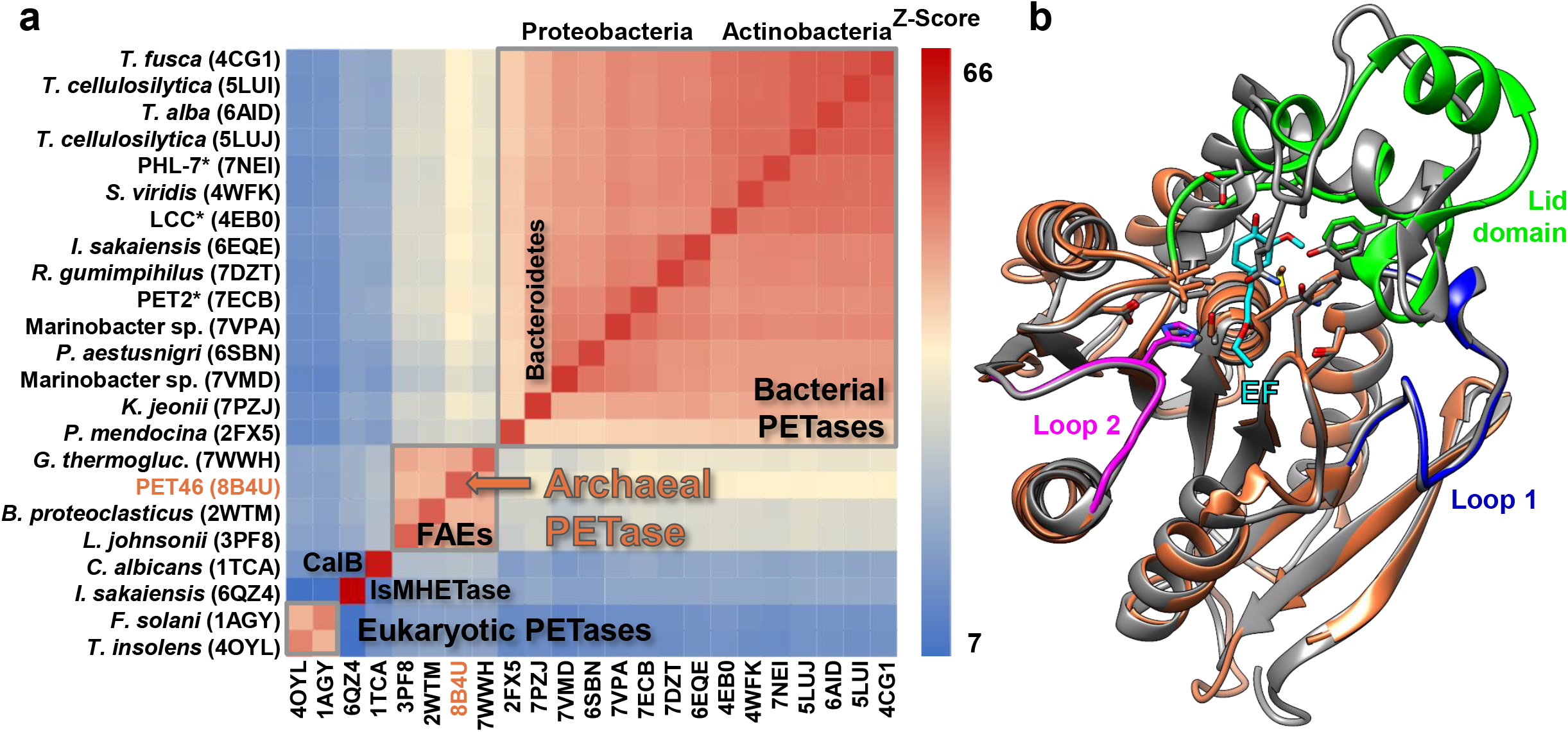
The protein structure of archaeal PETase PET46 and ferulic acid esterases (FAEs) is closely related to bacterial PETases. A heatmap represents structure similarity (Z-Score^69^) and reveals structural clusters. The FAE cluster, to which PET46 belongs, shows the highest similarity to the cluster of bacterial PETases. PET 46 is the FAE with the highest structural similarity to the bacterial PETases (**a**). PET46 shares most of its structure with FAEs (**b**). The structure of the archaeal PETase (coral orange) is overlaid to the crystal structure of the cinnamoyl esterase LJ0536 S106A mutant from *Lactobacillus johnsonii* (dark grey, PDB 3QM1) in complex with ethylferulate (EF, cyan). Loop 1 (deep blue) and Loop 2 (magenta) are highly conserved, but there are some variations in the lid domain (bright green). A Tyr in the loop of LJ0536 involved in substrate binding has a homologous Phe in PET46 (brilliant green). For structural alignments with other two FAEs and the tannase IsMHETase, see Supplementary Fig. S4. *No obvious phylogenetic affiliation.

PET46 and all three FAEs shared the highest structural similarity. Even Loop 1 and Loop 2 are highly conserved, but some variations are observed at the lid domain (Figure 3, Table 1 and Supplementary Fig. S4). PET46 and GthFAE share the “G-L-S-M-G” motif, very similar to the bacterial PETase’s “G-W/H-S-M-G”. The other two FAEs have “G-H-S-Q-G”, similar to eukaryotic PETase’s “G-Y-S-Q-G”. We analyzed the crystal structures of Est1E and LJ0536 co-crystallized with ferulic acid (FA; PDB 2WTN) or ethyl-ferulate (EF; PDB 3QM1) and confirmed that up to 5 aa in the lid are involved in substrate binding, including aromatic Tyr or Trp residues^38,39^, some of which PET46 also possesses (Figure 3, Supplementary Figs. S2 and S4). Overall, PET46 and FAEs build a cluster that is most similar to the cluster formed by bacterial PETases (Figure 3). The archaeal PETase is structurally most similar to the metagenomic bacterial PETase LipIAF5-2 (PET2^23^, Table 1). We then proceeded to characterize PET46 biochemically to confirm PETase activity.

### PET46 is a promiscuous feruloyl esterase that hydrolyzes MHET, BHET, 3PET and PET polymers

We tested PET46 for its activities on bis-(2-hydroxyethyl) terephthalate (BHET) and mono-(2-hydroxyethyl) terephthalate (MHET). Subsequently, we assayed activities on PET trimer (3PET) and polymers, both powder and foil. This clearly demonstrated that PET46 is able to break down plastic PET as well as the not completely hydrolyzed degradation products.

PET46 WT can degrade both BHET and MHET. In less than 30 min, all BHET (150 μM in 200 μL) was converted to MHET or terephthalic acid (TPA) in a 4:1 ratio. After 1 h of incubation, 51 μM TPA was measured. Incubation with the same amount of MHET for 1 h resulted in 52 μM TPA released (Figure 4). This implies that PET46 degrades BHET extremely efficiently, while MHET degradation occurred at the maximum rate independent from the starting substrate. PET46Δlid could degrade BHET to MHET, but we did not detect MHET degradation within 1 h incubation (Figure 4). Thus, the lid may be involved in substrate binding and catalysis.

**Figure 4:**
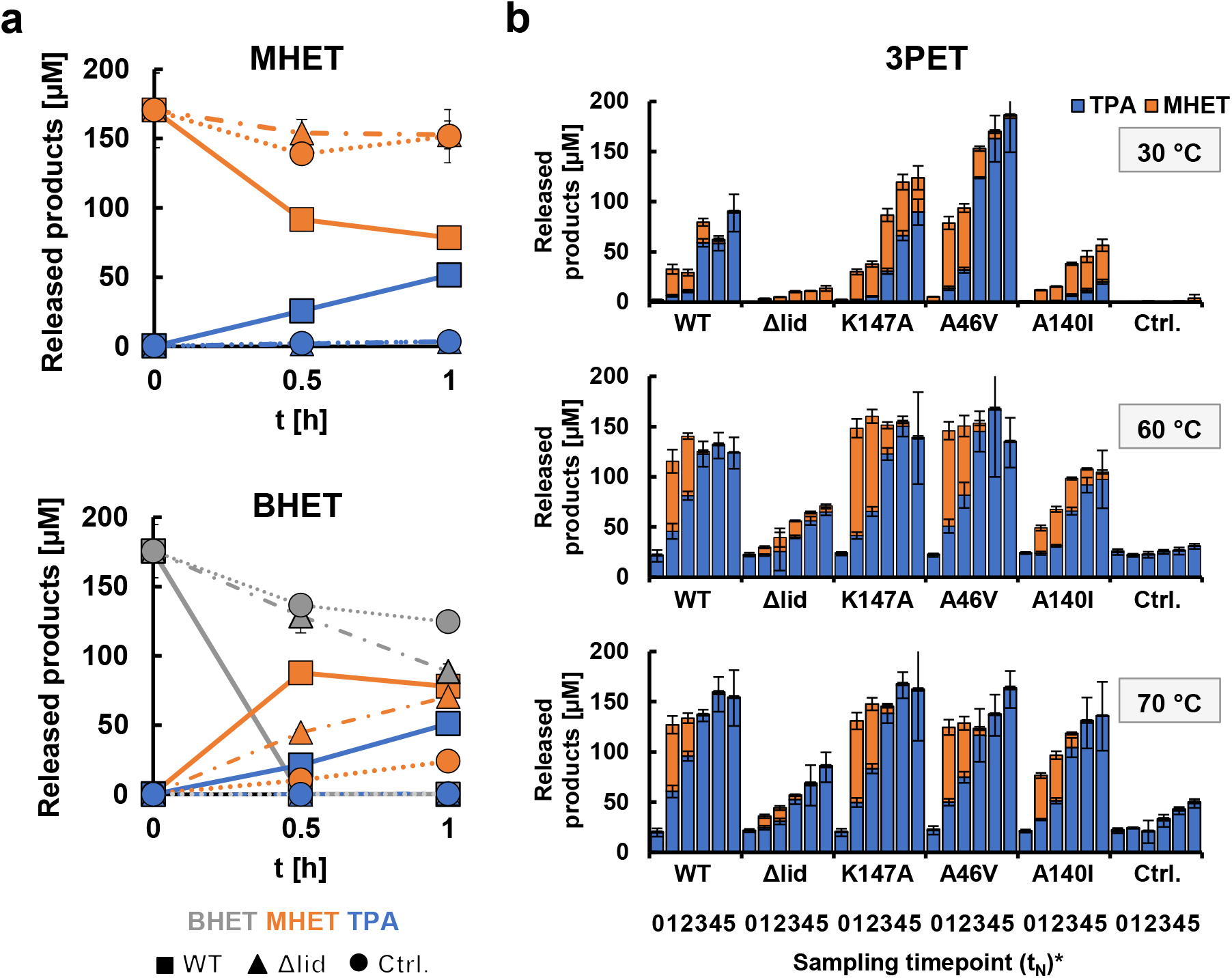
PET46 uses the lid domain to effectively degrade MHET, BHET and 3PET. PET46 WT can degrade both BHET and MHET to TPA and EG at 70 °C, but the lid-less variant PET46Δlid can only convert BHET to MHET (**a**). PET46 and the produced variants −1 degrade 3PET at 30, 60 and 70 °C (**b**). All experiments contain a total of 0.1 mg mL PET46 and 150 μM TPA equivalents in 200 μL potassium phosphate buffer pH 8. Error bars indicate the standard deviation of at least three replicates. *t_0_=0 h; t_1_=3 h; t_2_=6 h; t_3_=24 h; t_4_=48 h; t =72 h.

To better understand the BHET-degrading ability of PET46, its crystal structure was applied in docking experiments with this substrate. Two main clusters of docked poses were obtained, covering 83% and 12% of all solutions, and two smaller clusters containing 4% and 1% (Supplementary Fig. S5). For both main clusters, the smallest distance between the substrate’s carbonyl carbon and the hydroxyl oxygen from the catalytic serine is below 3.1 Å, indicating a plausible orientation of the substrate’s ester group towards the catalytic nucleophile (Supplementary Fig. S5). Based on the docking results, we identified two amino acids, A46 and A140, nearby both predominant docking poses that might be relevant for the substrate accessibility and binding (Supplementary Fig. S5). Introducing the larger substitutions A46V and A140I should thus impact the catalytic activity. We further identified K147, which possibly interacts with docked poses from the second-largest cluster. Variant K147A abolishes this interaction and widens the binding groove (Supplementary Fig. S5).

We then proceeded to incubate PET46 WT and all the constructed variants (including the PET46Δlid) on 3PET at 30, 60 and 70 °C. At the two highest temperatures, we observed a very similar activity pattern, where PET46 WT, K147A, and A46V degraded all the 3PET to MHET and TPA within the first 3 h (Figure 4). PET46 A140I performed slightly worse, while PET46Δlid could only convert half of the 3PET after 72 h incubation (Figure 4). Interestingly, A46V showed twice as much activity at 30 °C than the WT enzyme. In all experiments, we were not able to detect any BHET. Together with the previously obtained MHET-TPA profiles over time, we assume degradation happens at the polymer chain end (exo-activity), where 3PET is hydrolyzed to MHET units, which are subsequently converted to TPA and ethylene glycol (EG).

Finally, we assayed all PET variants on PET powder and film. The WT showed the highest activity of all enzymes and preferred PET powder over foil (Figure 5). We measured up to 62.38 μM TPA in 200 μL after one day at 70 °C from PET powder. On foil, a maximum of 4.45 μM TPA was released. The variants K147A, A46V, and A140I displayed only 45-50% less activity on powder than the WT, releasing 32.19-34.58 μM TPA equivalents from PET powder. On foil, they performed comparable to the WT. Finally, the lid-less variant released the lowest concentration of products regardless of the substrate, displaying up to 90% less activity on PET powder compared to the WT. As for the incubation with 3PET, we did not detect any BHET. Interestingly, after incubation with PET46 WT and A46V, no MHET was measured. This suggest that these two enzymes are more effective in its degradation than the other variants.

**Figure 5:**
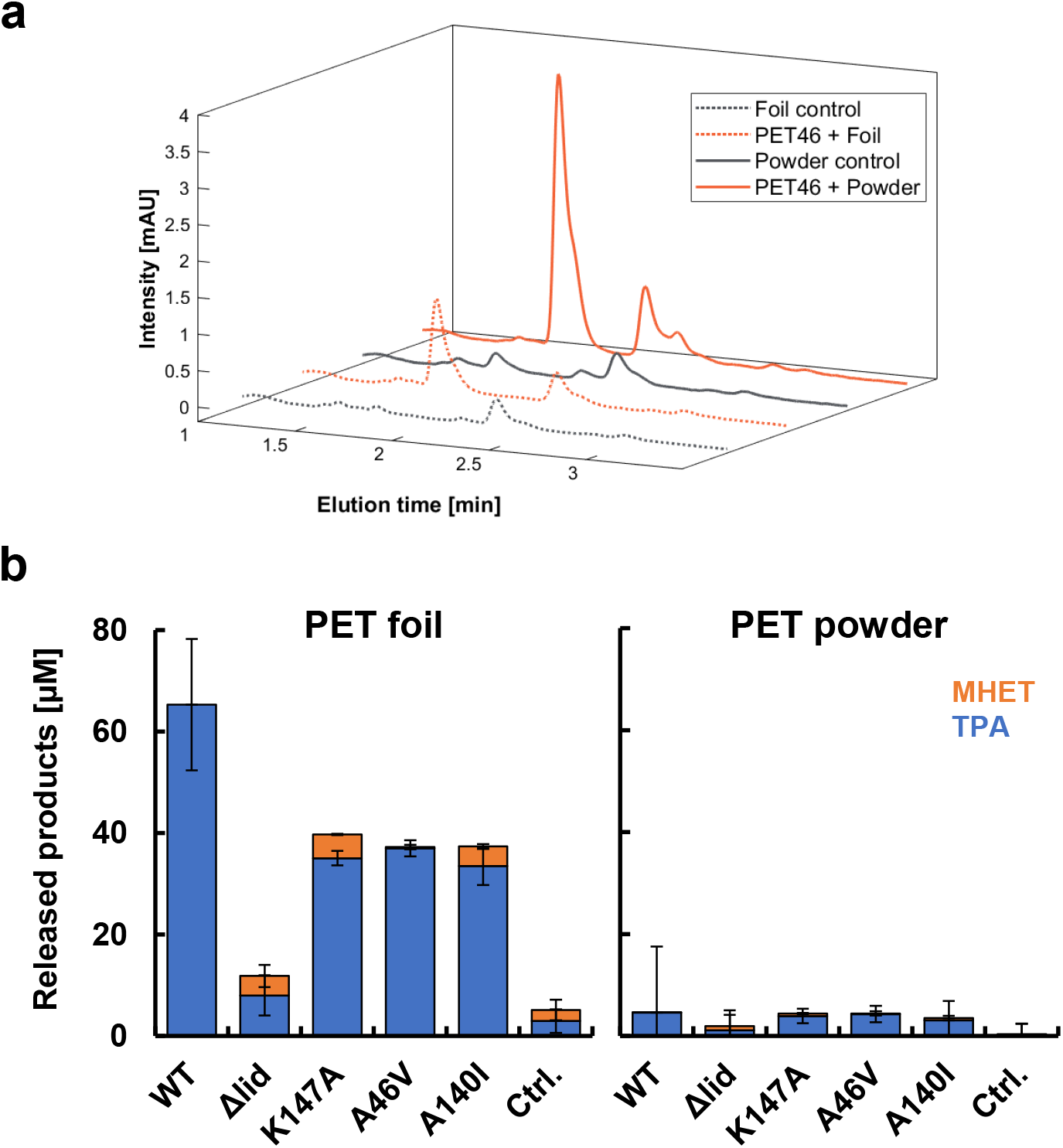
PET46 degrades PET polymer. UPLC chromatograms reveal a TPA peak (1.7 min) when incubating both PET powder and foil with PET46 WT for 24 h at 70 °C (**a**). 0.5 mg mL PET46 release up to 62 μM TPA out of PET powder and 4.5 μM out of PET foil after 24 h at 70 °C (**b**). No BHET could be measured. Data represent mean results from at least 3 replicates (3<n<5). Error bars indicate standard deviation.

We compared the measured activities of PET46 on PET substrates to literature values of the best-performing PET-active enzymes LCC and IsPETase. PET46 released 0.0052 μmol TPA mg^−1^ mL^−1^ h^−1^. Under optimal conditions, IsPETase^40^ releases 0.26-0.79 μmol TPA mg^−1^ mL^−1^ h^−1^. This makes PET46 50- to 150-fold less active, respectively, according to the literature values. However, an activity on PET polymer is clearly evident for PET46, which is higher than the one observed *e.g.* for the Bacteroidetes-derived PET30 enzyme^27^ (0.0003-0.0016 μmol TPA mg^−1^ mL^−1^ h^−1^). Furthermore, our work is the first report on PET degradation by a FAE.

Bathyarchaeota are ubiquitous and the predominant archaea at deep-sea environments like the Guaymas Basin^33,41^ and they have been shown to grow on lignin as energy source^42^, for which enzymes like PET46 need to be secreted. Thus, FAE-mediated promiscuous degradation of PET litter in the deep-sea seems plausible, even if at low rates.

### PET46 is adapted to the geochemical conditions at the Guaymas Basin

We characterized PET46 in more detail and with respect to its temperature and substrate profile. Therefore, a substrate spectrum was recorded with *p*NP-esters, which had an acyl chain length of 4 to 18 C-atoms.

The highest activities of PET46 were observed with *p*NP-decanoate (C10). PET46 was only poorly active on short (C4-C6) and long (C12-16) acyl chain lengths (Supplementary Fig. S6). The kinetic parameters for PET46 were determined with *p*NP-C10 at 70 °C and pH 8 according to Michaelis-Menten. Thereby, we measured a *V_max_* of 2.89*10^−5^ mol min^−1^, a *k*_cat_ of 110.99 min^−1^, a *K*_m_ of 0.4 mM and a *k*_cat_/*K*_m_ value of 277,475 M^−1^ min^−1^.

Using 1 mM *p*NP-decanoate as substrate, the recombinant enzyme PET46 revealed a relatively broad temperature spectrum. The highest activity was observed at 70 °C, while at 90 °C only 10% residual activity was detectable. The enzyme remained active at a temperature below 40 °C, but only had low activities (Figure 6). To further assess thermostability, the recombinant PET46 was incubated at 60 °C and 70 °C for two weeks. At 60 °C, the enzyme kept more than 60% of its activity for up to 8 days. At 70 °C, 80% of the activity was lost after 2 days, with only 10% remaining after 3 days (Figure 6). The original metagenomic sample was collected at a temperature of 48 °C^31^, at which PET46 shows 52% relative activity under laboratory conditions.

**Figure 6:**
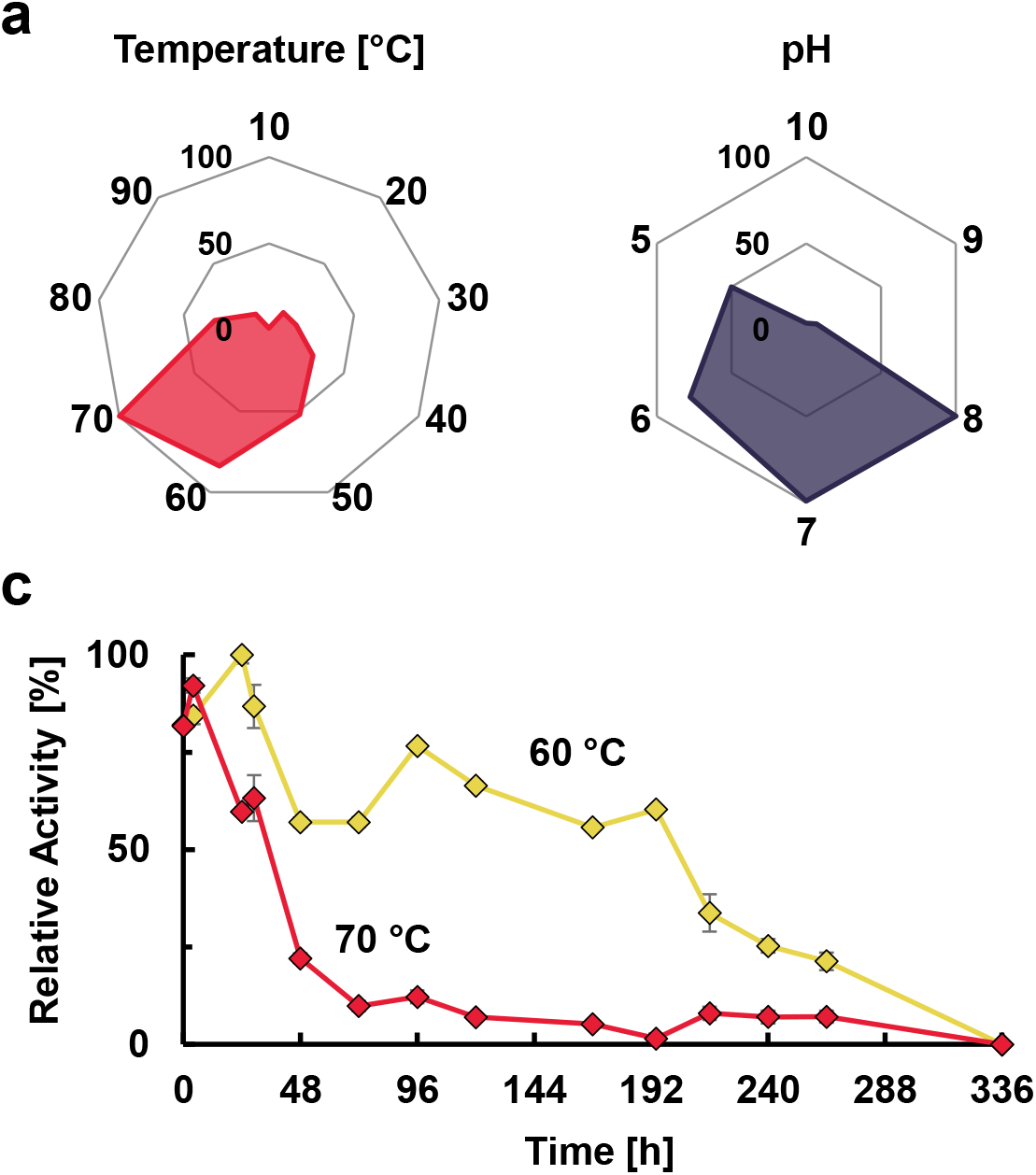
PET46 is a thermostable hydrolase adapted to the Guaymas Basin conditions. The enzyme’s optimal temperature and pH were determined by incubation with *p*NP-ester substrates (decanoate, C10) (**a**). The enzyme conserved most of its activity after 8-day incubation at 60 °C (**b**). Error bars indicate the standard deviation of at least three replicates. Standard deviation in (a) was below 6 % for all conditions assayed.

PET46 revealed activity for the broad pH range of 5-8. It had its optimum at pH 7-8 when tested on 1 mM *p*NP-C10 at 70 °C. However, it also retained relatively high activities (50%) at pH 5. The pH at the Guaymas Basin is recorded to be approximately 5.9^43^, at which PET46 would exhibit up to 77% of its activity under laboratory conditions.

To further characterize the effects of various metal ions, PET46 was incubated for 1 h with 1 or 10 mM Ca^2+^, Co^2+^, Cu^2+^, Fe^3+^, Mg^2+^, Mn^2+^, Ni^2+^ or Zn^2+^. The activity was assayed under optimal conditions and compared to a metal-free control. The activity of PET46 significantly increased in the presence of most of these ions. In contrast, Cu^2+^ reduced the activity by 50%. Especially the addition of Zn^2+^ resulted in almost two-fold activity increase (Supplementary Fig. S6). Some of these ions are present at significant concentrations in the Guaymas Basin^44^. Thus, metal binding to the protein seems plausible.

Further, we tested the sensitivity of PET46 towards detergents and the reducing agent DTT. A concentration of 1 and 5% of the detergents Triton X-100 and DTT strongly affected the enzyme activities (Supplementary Fig. S6). Interestingly, 1% DTT stimulated esterase activity by a factor of two.

Finally, we assayed the solvent tolerance of PET46. In general, the enzyme was remarkably stable in the presence of acetone, DMF, isopropanol, and DMSO. Notably, 10% acetone and 5% DMSO and DMF increased the enzyme’s activities by a factor of 2 (Supplementary Fig. S6). This is a noteworthy solvent tolerance, which makes it an ideal candidate for future biotechnological applications (*e.g.* in a multi-enzyme PET degradation approach^28^).

Overall, PET46 is a well-adapted and very stable enzyme in its natural environment. Together with our results on PET poly-oligo- and monomers hydrolysis, we conclude that enzymes associated with lignin degradation, and especially FAEs from Bathyarchaeota and other prokaryotes, may have a global impact in promiscuity-driven degradation of PET litter in the deep-ocean.

## DISCUSSION

Plastic pollution is now considered one of the world’s greatest threats to the environment and global health. Among different plastics, PET is discharged in large quantities into the environment where it accumulates. Our knowledge of microbial degradation processes in nature is very sparse. Since PET is composed of ester bonds that can be hydrolyzed by enzymes, a significant number of bacterial and a few fungal genes encoding those have been identified in previous research.

PET-degrading enzymes belong to the classes of cutinases [EC (enzyme category) 3.1.1.74], lipases (EC 3.1.1.3), or carboxylesterases (EC 3.1.1.1), and these can only hydrolyze amorphous and low-crystalline PET. The PET-active enzymes hydrolyze the ester bond to produce either BHET, MHET, or TPA and EG.

Many of the PET-active enzymes are thermostable and perform best at temperatures between 55 and 65 °C. This temperature is close to the glass transition temperature of PET (65-71 °C) and favors the formation of softer, more flexible domains with better accessibility for the enzymes. However, all known native PET-active hydrolases have a rather low catalytic activity towards high molecular weight PET and all are promiscuous enzymes, implying that PET is not the native substrate. Notably, esterases are well known to be promiscuous enzymes. Some of these enzymes are known to turn over more than 70 different chemical substrates^45,46^.

Current research on this topic mainly follows two major goals. First, much research is directed to the design and evolution of efficient catalysts for recycling of PET. The second focus lies in mining biodiversity to better understand their roles in nature, global distribution patterns and obtain novel enzymes with improved traits that can be used as backbones for better catalysts. Our study aimed to unravel novel structural and phylogenetic biodiversity of PET-degrading enzymes.

Within this setting, we provide strong evidence that the Candidatus Bathyarchaeota archaeon MAG hosts the promiscuous esterase PET46 that can act on amorphous and low-crystalline PET. Our data imply that PET46 has PETase activity when incubated with PET powder. Besides, PET46 hydrolyzes BHET and MHET with significant rates, confirming that it can handle both the polymer and the intermediates (Figures 4 and 5).

Based on its structural analysis, PET46 is a feruloyl esterase. Feruloyl esterases (FAE; EC 3.1.1.73) release ferulic acid and other hydroxycinnamic acids from plant-based hemicellulose and lignin, which has a large biotechnological application^47^. They are widespread in nature and have been found in bacteria, plants, and fungi. Notably, this is the first report on an active and functionally verified archaeal FAE. Their 3D structure usually reveals a canonical eight-strand α/β-fold of lipases and esterases. In addition, a lid domain is observed, which, analogous to lipases, confines the active site with a loop that confers plasticity to the substrate-binding site^48^.

With respect to the degradation of PET, none of the currently known PETases is a FAE. A recent study described a metagenomic FAE with phthalate-degrading activity, but no PET degradation was assayed^49^. PETases are assumed not to have a lid domain. However, some enzymes acting on the intermediate MHET are annotated as tannases, which form a protein family with FAEs, and they bear a lid domain of varying length. They hydrolyze MHET into TPA and EG. One of the best-studied examples is the MHETase derived from the gram-negative bacterium *Ideonella sakaiensis.* This enzyme acts on MHET, but recently an exo-function on PET pentamers was described^50^. It also hydrolyzed BHET in a concentration-dependent manner, and its three-dimensional structure shows a much larger lid domain involving more than 200 aa (Supplementary Fig. S4). This marked structural difference may possibly explain the difference in the substrate specificities.

PET46 is encoded in a marine Bathyarcheota MAG. Microorganisms affiliated with the Bathyarchaeota are globally occurring and widespread in marine and terrestrial anoxic sediments^51^. They can use a wide range of polymers as carbon and energy source, and they are well known to be very versatile with respect to the metabolic capabilities. They are further known to be relatively abundant in some marine sediments. Because of their huge metabolic potential, it is further assumed that they may play a significant role in global carbon biogeochemical cycling^51–54^. Interestingly, Bathyarcheota have been associated with the degradation of the biopolymer lignocellulose previously^42^. Therefore, the observation here that not only their genomes code for FAEs, but also the demonstration that they are functionally active underscores this observation.

Within this setting, our observations that PET46 was catalytically active on PET powder is in line with the known wide metabolic diversity of the Bathycharchaeota^55^. Nevertheless, when we benchmarked our enzyme with the well-characterized enzyme IsPETase, our data implied that the overall PETase activities observed for PET46 are significantly lower, but were higher than those published for the Bacteroidetes-derived enzyme PET30. Nevertheless, it is noteworthy that PET-activities can hardly be compared between studies from different laboratories, as many influencing factors like sample preparation and purity, assay conditions, and measurement methods strongly vary.

While PET esterases are not highly conserved among each other, few structural traits and sequence homologies appear to be common in most of the known enzymes (Figure 3). Based on our data analyses and others^30^, it becomes evident that none of the current enzymes carries a lid domain, which showed to be crucial for enzymatic activity of PET46. All published active enzymes are secreted proteins, that carry at least an N-terminal signal peptide and some even a PorC-like type 9 secretion system (T9SS)-affiliated signal^27^. The region involved in substrate binding contains in general the amino acids Tyr/Phe-Met-Trp/Tyr, and the catalytic triad is composed of Asp-His-Ser. Further, active enzymes carry 1-2 disulfide bonds and of these, one is close to the active site. The active site is well accessible for the bulky substrates and is located in a rather large cavity. For more detailed analyses of common PETase features, we refer to other studies^13,30^.

In summary, our biochemical results significantly extend the knowledge of PETase enzymes and their biodiversity. Our study further enables the development of an expanded phylogenetic framework for identifying the diversity of putative PETases in marine microbial groups throughout the global ocean. Finally, the data presented here will help to advance our knowledge on the ecological role of the Bathyarchaeota and with respect to the possible decomposition of marine PET litter.

## METHODS

### Profile Hidden-Markov Model (HMM) searches identify putative archaeal PETases

An HMM constructed from all PET-degrading enzymes listed in the PAZy database^16^ was used to search against NCBI’s non-redundant protein database (ftp.ncbi.nlm.nih.gov/blast/db/FASTA/nr.gz) filtered for sequences of archaeal origin (tax ID: 2157) as described previously^13,23,32^.

### Primers, constructs and bacterial strains used

The gene coding for PET46 was codon-optimized and synthesized in pET21a(+) (Novogene, Cambridge, UK) by Biomatik (Ontario, Canada) and transformed in *Escherichia coli* BL21 (DE3) (Novagen/Merck, Darmstadt, Germany) for protein production. The primers used to generate all PET46 mutants by site directed mutagenesis were synthesized by Eurofins Genomics (Ebersberg, Germany), and are listed in Supplementary Table S2. Sequencing of all constructs was conducted by Mycrosynth Seqlab GmbH (Göttingen, Germany).

### Protein production

PET46 WT and its mutant derivatives were produced heterologously by growing *E. coli* BL21 (DE3) cells carrying the respective pET21a(+) construct at 37 °C in Luria-Bertani (LB) medium containing 100 μg mL^−1^ ampicillin. When OD_600_ reached 0.7, 1 mM IPTG was added to induce expression of the genes and cultures were incubated overnight at 22 °C to facilitate protein production. Cells were centrifuged and lysis was carried out via French Press three times (1,250 psi). The proteins were purified from the cleared lysate with Ni-NTA agarose (Macherey-Nagel, Düren, Germany) following concentration and dialysis against 0.1 M potassium phosphate buffer pH 7.

### Crystallization data collection, data reduction, structure determination, refinement and final model analysis

PET46 was crystallized by sitting-drop vapor-diffusion at 12 °C at a concentration of 10 mg mL^−1^ in 100 mM potassium phosphate buffer pH 7. 1.5 μL of PET46 were mixed with 1.5 μL of reservoir solution consisting of 325 mM (NH_4_)H_2_PO_4_. Crystals formed after 3-4 weeks, were harvested and cryo-protected with 35% ethylene glycol followed by flash-freezing in liquid nitrogen. Diffraction data were collected at −173 °C (100 K) at beamline ID23-1 (ESRF, Grenoble, France) using a 0.9793 Å wavelength. Data reduction was performed using XDS^56^ and Aimless^57^ from the CCP4 Suite^58^. The structure was solved via molecular replacement with Phaser^59^ using an AlphaFold^60^ model as search model. The initial model was refined alternating cycles of manual model building in COOT^61,62^ and automatic refinement using Phenix^63^ v.1.19.2_4158. Data collection and refinement statistics are reported in Supplementary Table S1. The structure assembly was analyzed using PISA^64^.

### Sequence and structure analysis

Local alignments were performed with BLASTp^65^ or DIAMOND^66^ v.2.0.15, and network analysis was carried out in Cytoscape^67^ v.3.9.1. Conserved domains at the sequence level were inferred from the Conserved Domain Database^68^ (CDD). Heuristic structural searches against the Protein Databank (PDB) were performed on the Dali server^69^. Structural visualization and alignments were performed with PyMOL^70^ v.2.0 and USFC Chimera^71^ v.1.16.

### Substrate docking

The BHET substrate was docked into the catalytic site of PET46 utilizing a combination of AutoDock3^72^ as a docking engine and DrugScore2018^73,74^ as an objective function. Following an established procedure^73,75^, the docking protocol considered 100 independent runs for BHET using an initial population size of 150 individuals, a maximum number of 50.0 × 10^3^ generations, a maximum number of 1.0 × 10^6^ energy evaluations, a mutation rate of 0.02, a crossover rate of 0.8, and an elitism value of 1. The Lamarckian genetic algorithm was chosen for sampling in all approaches. The distance between the carbonyl carbon from the docked BHET and the hydroxyl oxygen from the catalytic serine was measured using the PyMOL Molecular Graphics System^70^ v.2.3.0.

### PET degradation assays

Respectively, 0.1 mg mL^−1^ PET46 WT and the generated variants were incubated with 50 μM ethylene terephthalate linear trimer (3PET, Toronto Research Chemicals, Ontario, Canada), 150 μM bis-(2-hydroxyethyl) terephthalate (BHET), 150 μM mono-(2-hydroxyethyl) terephthalate (MHET; Merck, Darmstadt, Germany). Alternatively, 0.5 mg mL^−1^ enzyme were incubated with 7 mg PET foil platelet (a=5 mm^2^, 33.6 μmol or 168 mM TPA eq.), or 2 mg semi-crystalline PET powder (9.6 μmol or 48 mM TPA eq.; GoodFellow GmbH, Hamburg, Germany). The reaction took place in 200 μL with 0.1 M potassium-phosphate buffer pH 8 at 30, 60, or 70 °C for a maximum of 5 days. For end point analysis, samples were prepared in 96 well microtiter plates by adding 12.5 μL reaction supernatant to 50 μL acetonitrile with 1% v/v trifluoroacetic acid (TFA) followed by centrifugation (2,204 *g*, 30 min; A-2-DWP rotor, Eppendorf AG, Hamburg, Germany) and transferring of 50 μL centrifugation supernatant into 150 μL MilliQ H_2_O. Samples were sealed using ZoneFree™ sealing film (Excel Scientific, Victorville, CA, USA) and stored at −20 °C until analysis. Samples were analyzed via RP-UPLC as described previously^27^. Standards of the expected degradation products TPA, MHET, and BHET were analyzed to obtain the respective elution times. All assays were performed in triplicates and compared to an enzyme-free control.

### Biochemical characterization

Initial biochemical characterization aimed to identify the WT enzyme’s optimal temperature, pH, and substrate chain length and was performed with *para*-nitrophenyl (*p*NP) esters as described previously^23,27^. To test the thermostability of the enzyme, it was incubated at 60 and 70 °C for up to two weeks prior to a *p*NP assay under optimal conditions to quantify residual activity. Furthermore, the effect of metal ions, detergents, and organic solvents was assayed. The enzyme was either pre-incubated for one hour with 1 or 10 mM Ca^2+^, Co^2+^, Cu^2+^, Fe^3+^, Mg^2+^, Mn^2+^, Ni^2+^ or Zn^2+^ (chloride salts) or different detergents and organic solvents were added to the standard reaction mixture (Supplementary Fig. S5).

To test for general ferulic acid esterase activity, a colorimetric pH-shift-based assay with the model substrate ethyl cinnamate (EC), and was performed as described previously^46^. In short, the reactions took place in 5 mM EPPS buffer with 0.45 mM phenol red. The release of protons due to enzymatic cleavage of the ester results in a decrease in absorbance at 550 nm, which is measured photometrically.

All assays were performed in triplicates and compared to an enzyme- or additive-free control.

## ACKNOWLEDGEMENTS

X-ray diffraction measurements were performed on beamline ID23-1-3 at the European Synchrotron Radiation Facility (ESRF), Grenoble, France. This work was in part supported by the European Commission (Horizon2020 project FuturEnzyme; grant agreement ID 101000327) and the Federal Ministry of Education and Research (BMBF) within the programs MarBiotech (031B0562A), MetagenLig (031B0571A and 031B0571B), LipoBiocat (031B0837B) and PlastiSea (031B867B and 031B867F) at the Universities of Hamburg and Kiel and LipoBiocat (031B0837A) at the HHU Düsseldorf. The Center for Structural Studies is funded by the Deutsche Forschungsgemeinschaft (DFG Grant number 417919780 and INST 208/740-1 FUGG). HG is grateful for computational support and infrastructure provided by the ‘‘Zentrum für Informations- und Medientechnologie” (ZIM) at the Heinrich Heine University Düsseldorf and the computing time provided by the John von Neumann Institute for Computing (NIC) on the supercomputer JUWELS at Jülich Supercomputing Centre (JSC) (user ID: HKF7, VSK33, lipases).

## AUTHOR CONTRIBUTIONS

P.P.G., J.C., R.A.S. and W.R.S designed the study. P.P.G., M.F.G., G.F. and P.T. were involved in enzyme production, mutation experiments and biochemical assays. M.F.G. and R.F.D. executed UPLC analysis. D.D. performed HMM searches. V.A., E.C., J.S. and S.H.S. performed crystallization experiments and structure solving. P.P.G. performed structural analysis. C.P., J.D. and H.G. were involved in ligand docking. W.R.S., R.A.S., S.H.S. and H.G. received funding. P.P.G., J.C. and W.R.S. wrote the first draft of the manuscript with input from all authors.

## SUPPLEMENTARY FIGURE LEGENDS

**Supplementary Fig. S1: The gene coding for PET46 is inserted between genes related with translation and has bacterial homologs.** PET46 is located in a small contig between genes coding for translation-associated proteins. It contains conserved sequence domains from dipeptidyl aminopeptidase/acylaminoacyl peptidase (DAP2), acetyl xylan esterase (AXE1), dienelactone hydrolase (DLH) or lysophospholipase (PldB, **a**). Archaeal homologs from PET46 derive mainly from other Bathyarchaeota, but there are more bacterial homologs (query cov. > 80%, seq. id > 40 %) (**b**). A sequence network analysis displaying sequence similarity (bit score) reveals that PET46 and its archaeal homologs share high homology to the Firmicutes and Planctomycetes sequences (**c**). Archaeal sequences are displayed as circles and bacterial as triangles. PET46 is highlighted with a yellow border. Color legend is shared with “b”. The most abundant phylum within a group is showed in parenthesis.

**Supplementary Fig. S2: The crystal structure of PET46 consists of 7 o-helixes and 8 β-strands forming the canonical α/β-hydrolase fold and 3 o-helixes and 2 anti-parallel β-strands making the lid.** Together with the lid domain (bright green), Loop 1 and Loop 2 (deep blue and magenta) are the main structural variations with the IsPETase (**a**). These loops are conserved in all ferulic acid esterases (FAEs) analyzed. Displayed are the catalytic triad and homologous residues involved in substrate binding in PETases or FAEs. The lid domain contains at least two aromatic residues (Phe148 and Trp172; bright green). 2Fo-Fc map contoured at one sigma is shown as blue mesh around the PO_4_ and ethylene glycol (EG) moieties modelled near the active site (**b**). Stereo image of the density of the active site residues (**c**). The catalytic triad residues are shown as sticks.

**Supplementary Fig. S3: Hydroxycinnamic acid-esters, the native substrates of ferulic acid esterases, are similar to the terminus of a PET polymer.** A feruloyl-polysaccharide (left) and a *p*-coumaryl-polysaccharide (right) are two examples of hydroxycinnamic acid-polysaccharide esters (**a**). The synthetic ethylene terephthalate linear trimer (3PET) used as a substrate in this study (**b**). The attacked oxygen during an exo-reaction is highlighted with an arrow. PET46 degrades ethyl cinnamate (EC), a model substrate for FAE activity (**c**). A pH-shift assay (phenol red) with ethyl cinnamate (EC) and PET46 at two concentrations + results in the release of H upon ester hydrolysis.

**Supplementary Fig. S4: The archaeal PETase PET46 is structurally homolog to ferulic acid esterases (FAEs).** The crystal structure of PET46 (coral orange) is compared to the crystal structures of GthFAE from *Geobacillus thermoglucosidasius* (lime green; PDB 7WWH; **a**), the Est1E FAE from *Butyrivibrio proteoclasticus* (cream white; PDB 2WTN) bound to ferulic acid (FA; cyan; **b**), and the tannase IsMHETase from *I. sakaiensis* (petrol green; PDB 6QZ4; **c**). The lid domains of PET46 and IsMHETase have been omitted in “c” for better visualization (bright green).

**Supplementary Fig. S5: Docking of BHET into PET46**. Docking of BHET yielded four possible binding poses (clusters CL1-CL4) in PET46 (**a**). Docked poses of the two largest clusters within PET46 with the box depicting the search space (**b**). Distributions of the smallest distances between the docked substrate’s carbonyl carbon and the hydroxyl oxygen from the catalytic serine for the two largest clusters (**c**, **d**). Location of the substituted amino acids in the A46V variant (blue sticks, **e**), the A140I variant (blue sticks, **f**), and the K147 variant (white sticks, **g**) of PET46. The comparison of the substrate binding sites for the (H) WT (white surface, **h**) and the K147A variant (gray surface, **i**) shows an extended substrate binding site in the variant. The same orientation is used as for “g”.

**Supplementary Fig. S6: Biochemical characterization of PET46.** Optimal *p*NP-ester acyl chain length was determined (**a**). The effect of metal ions (**b**), detergents (**c**) and organic solvents (**d**) on the activity of PET46 was studied compared to an additive-free control (Ctrl.). Error bars indicate the standard deviation of at least three replicates. Standard deviation in “a“ was below 6 %.

## SUPPLEMENTARY TABLE LEGENDS

**Supplementary Table S1: Data collection and refinement statistics.** Values in parenthesis refer to the highest resolution shell.

**Supplementary Table S2: Primers used in this study.** Lid deletion and point mutations were introduced by site-directed mutagenesis. pET primers were used for Sanger sequencing to verify the correctness of the produced variants prior to expression.

