## Supplemental Tables for "The first archaeal PET-degrading enzyme belongs to the feruloyl-esterase family"

### SUPPLEMENTARY TABLES

**Supplementary Table S1: Data collection and refinement statistics.** Values in parenthesis refer to the highest resolution shell.

| **PET46** | |
|---|---|
| **PDB ID** | 8B4U |
| **Data collection** | |
| **Wavelength (Å)** | 0.9793 |
| **Space group** | $P6_122$ |
| ***Unit Cell Parameters*** | |
| **a, b, c (Å)** | 79.29 79.29 171.55 |
| **α, β, γ (°)** | 90.00 90.00 120.00 |
| **Resolution (Å)** | 68.67 - 1.71 (1.74 - 1.71) |
| **Number of unique reflections** | 34960 (1813) |
| **$R_{merge}$** | 0.066 (1.370) |
| **$R_{meas}$** | 0.071 (1.457) |
| **$R_{pim}$** | 0.024 (0.487) |
| **<I/σ(I)>** | 15.7 (1.5) |
| **$CC^{1/2}$** | 0.999 (0.559) |
| **Completeness (%)** | 99.4 (99.9) |
| **Multiplicity** | 8.5 (8.5) |
| **Refinement** | |
| **Resolution (Å)** | 38.68-1.71 |
| **Number of reflections** | 34953 |
| **$R_{work}$/$R_{free}$ (%)** | 15.23 / 17.27 |
| ***R.M.S. deviations*** | |
| **bond length (Å)** | 0.011 |
| **bond angles (°)** | 1.013 |
| ***Ramachandran plot*** | |
| **favored (%)** | 98.88 |
| **allowed (%)** | 1.12 |
| **outliers (%)** | 0 |

**Supplementary Table S2: Primers used in this study.** Lid deletion and point mutations were introduced by site-directed mutagenesis. pET primers were used for Sanger sequencing to verify the correctness of the produced variants prior to expression.

| Primer Name | Nucleotid Sequence (5‘->3‘) | $T_m$ [°C] | Reference |
|---|---|---|---|
| Δlid_for | TTCTACCTTAGATGTGCTGGATCGC | 67.1 | This work |
| Δlid_rev | GAGTCCCATGCACTATACAGAATAACAAATTTAATG | 67.3 | This work |
| K147A_for | AGCCTTAACCCCGCTGCGCCGCGCGTTTCTG | 81.9 | This work |
| K147A_rev | GACGAATTGATTCTTTTTCTAAGCCTTCCAGAAACGCGCGGCG | 77.7 | This work |
| A46V_for | TTTCATGGCTTTACCGGCAATAAATCAGAAGTGCATCGTCTGT | 76.0 | This work |
| A46V_rev | GCGTGCAACATGAACAAACAGACGATGCACTTCTGATTT | 75.2 | This work |
| A140I_for | TCGTCGCATTAAATTTGTTATTCTGTATAGTGCAATTTTAACCCCGC | 73.9 | This work |
| A140I_rev | ATTTGCGGCGCAGCGGGGTTAAAATTGCACTATAC | 75.9 | This work |
| pET_for | ATATAGGCGCCAGCAACC | 62.7 | Novagen/Merck (Darmstadt, Germany) |
| pET_rev | TCCGGATATAGTTCCTC | 54.3 | Novagen/Merck (Darmstadt, Germany) |
