## Supplemental Figures for "The first archaeal PET-degrading enzyme belongs to the feruloyl-esterase family"

### Supplementary Figures

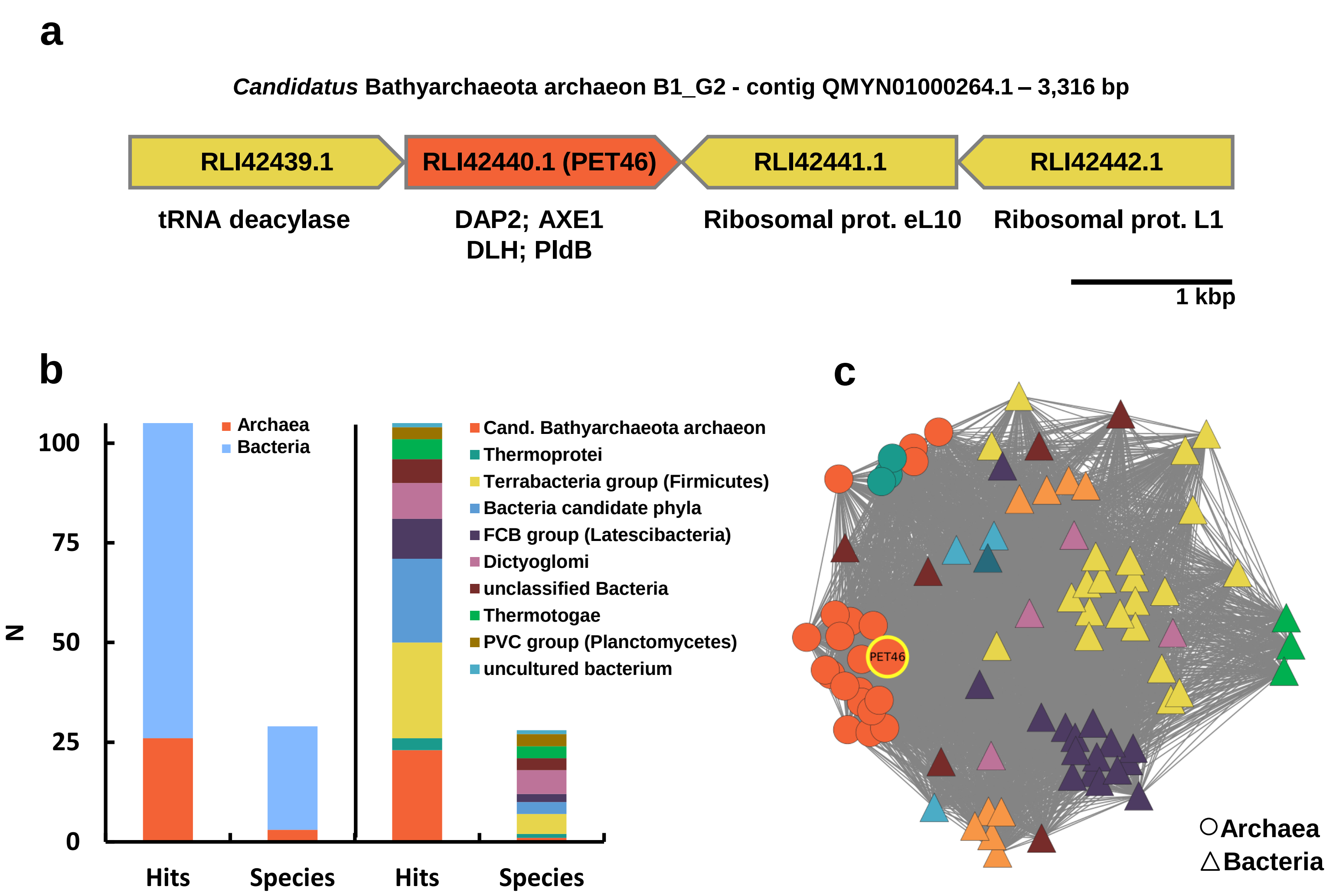


**Supplementary Fig. S1: The gene coding for PET46 is inserted between genes related with translation and has bacterial homologs.** PET46 is located in a small contig between genes coding for translation-associated proteins. It contains conserved sequence domains from dipeptidyl aminopeptidase/acylaminoacyl peptidase (DAP2), acetyl xylan esterase (AXE1), dienelactone hydrolase (DLH) or lysophospholipase (PldB, **a**). Archaeal homologs from PET46 derive mainly from other Bathyarchaeota, but there are more bacterial homologs (query cov. > 80%, seq. id > 40 %) (**b**). A sequence network analysis displaying sequence similarity (bit score) reveals that PET46 and its archaeal homologs share high homology to the Firmicutes and Planctomycetes sequences (**c**). Archaeal sequences are displayed as circles and bacterial as triangles. PET46 is highlighted with a yellow border. Color legend is shared with “b”. The most abundant phylum within a group is showed in parenthesis.

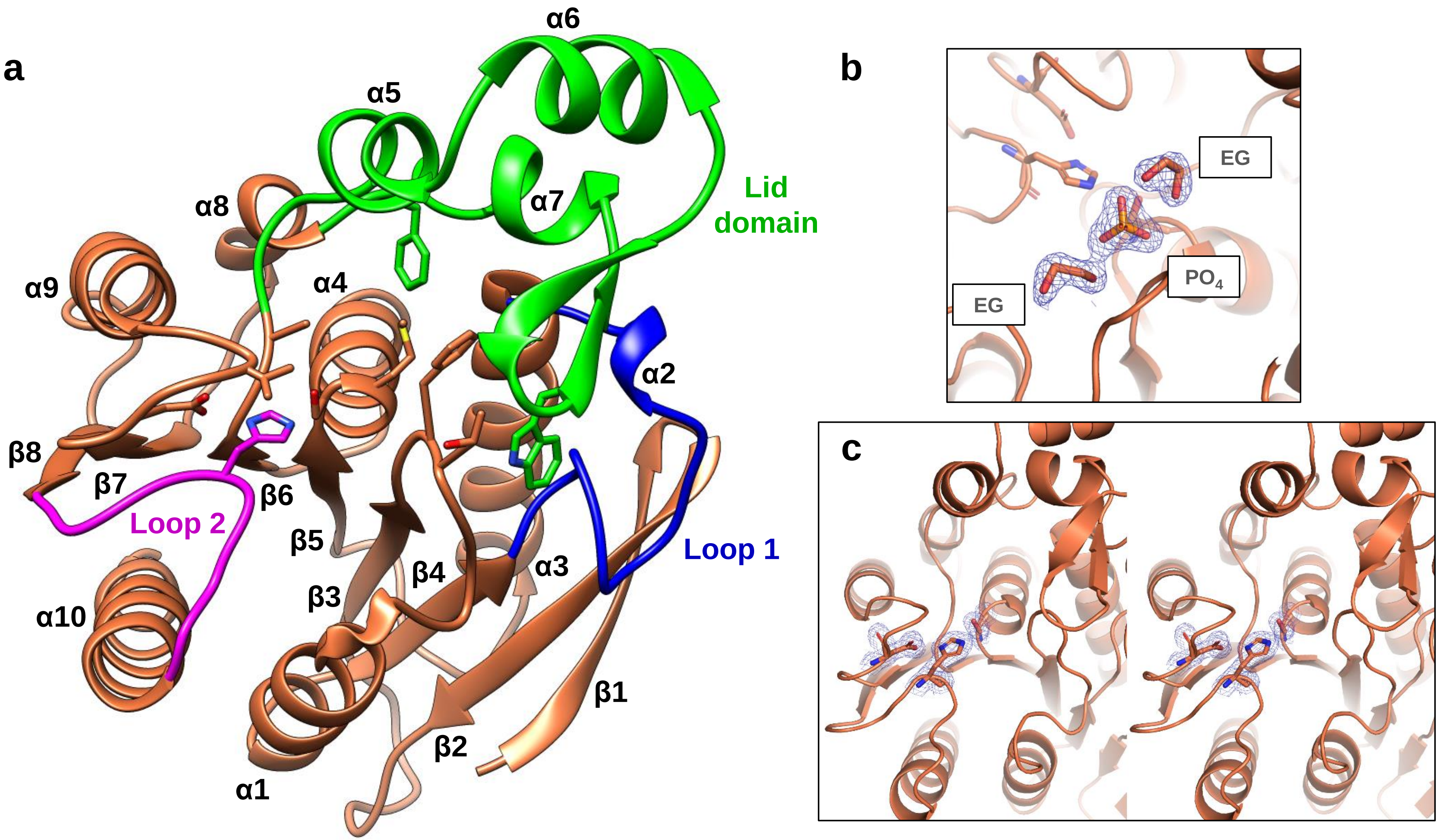


**Supplementary Fig. S2: The crystal structure of PET46 consists of 7 α-helixes and 8 β-strands forming the canonical α/β-hydrolase fold and 3 α-helixes and 2 anti-parallel β-strands making the lid.** Together with the lid domain (bright green), Loop 1 and Loop 2 (deep blue and magenta) are the main structural variations with the IsPETase (**a**). These loops are conserved in all ferulic acid esterases (FAEs) analyzed. Displayed are the catalytic triad and homologous residues involved in substrate binding in PETases or FAEs. The Lid Domain contains at least two aromatic residues (Phe148 and Trp172; bright green). 2Fo-Fc map contoured at one sigma is shown as blue mesh around the $PO_4$ and ethylene glycol (EG) moieties modelled near the active site (**b**). Stereo image of the density of the active site residues (**c**). The catalytic triad residues are shown as sticks.

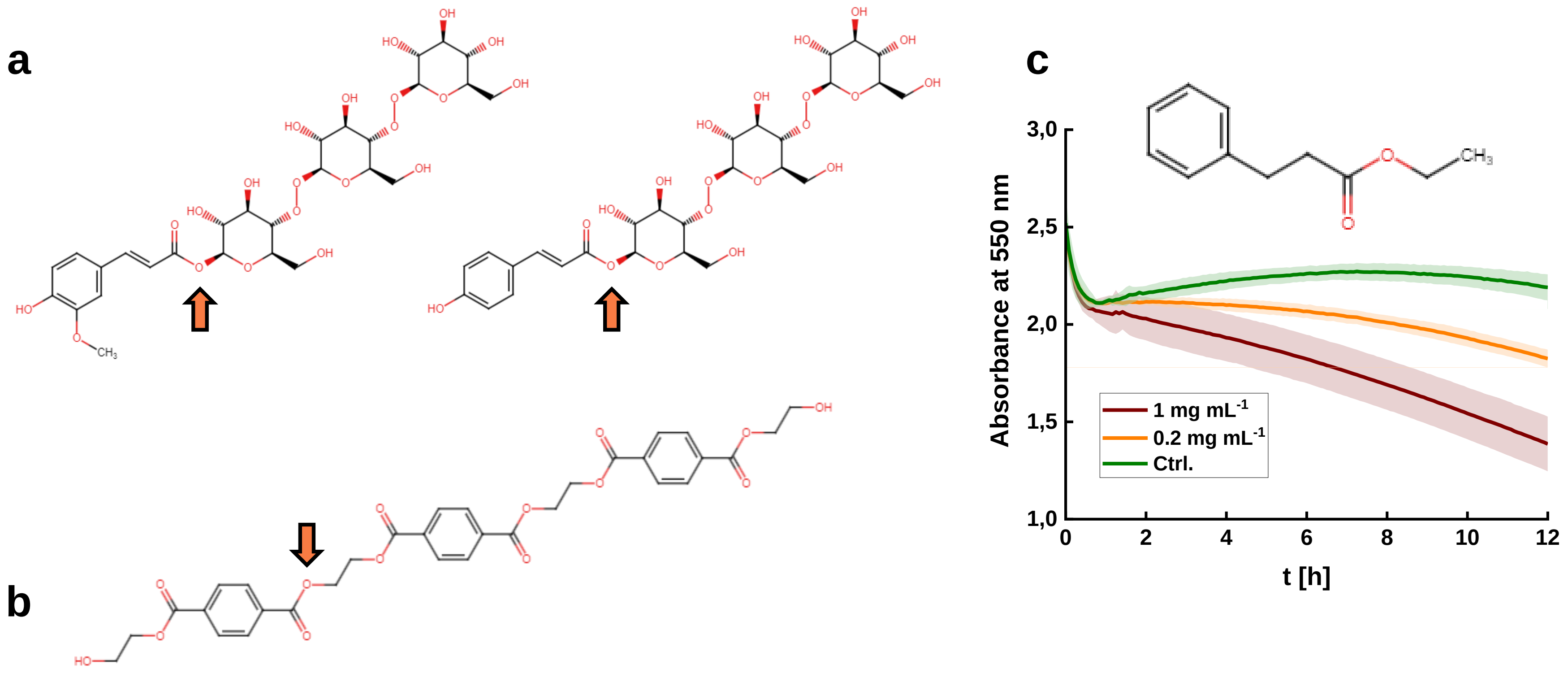


**Supplementary Fig. S3: Hydroxycinnamic acid-esters, the native substrates of ferulic acid esterases, are similar to the terminus of a PET polymer.** A feruloyl-polysaccharide (left) and a *p*-coumaryl-polysaccharide (right) are two examples of hydroxycinnamic acid-polysaccharide esters (**a**). The synthetic ethylene terephthalate linear trimer (3PET) used as a substrate in this study (**b**). The attacked oxygen during an exo-reaction is highlighted with an arrow. PET46 degrades ethyl cinnamate (EC), a model substrate for FAE activity (**c**). A pH-shift assay (phenol red) with ethyl cinnamate (EC) and PET46 at two concentrations results in the release of $H^+$ upon ester hydrolysis.

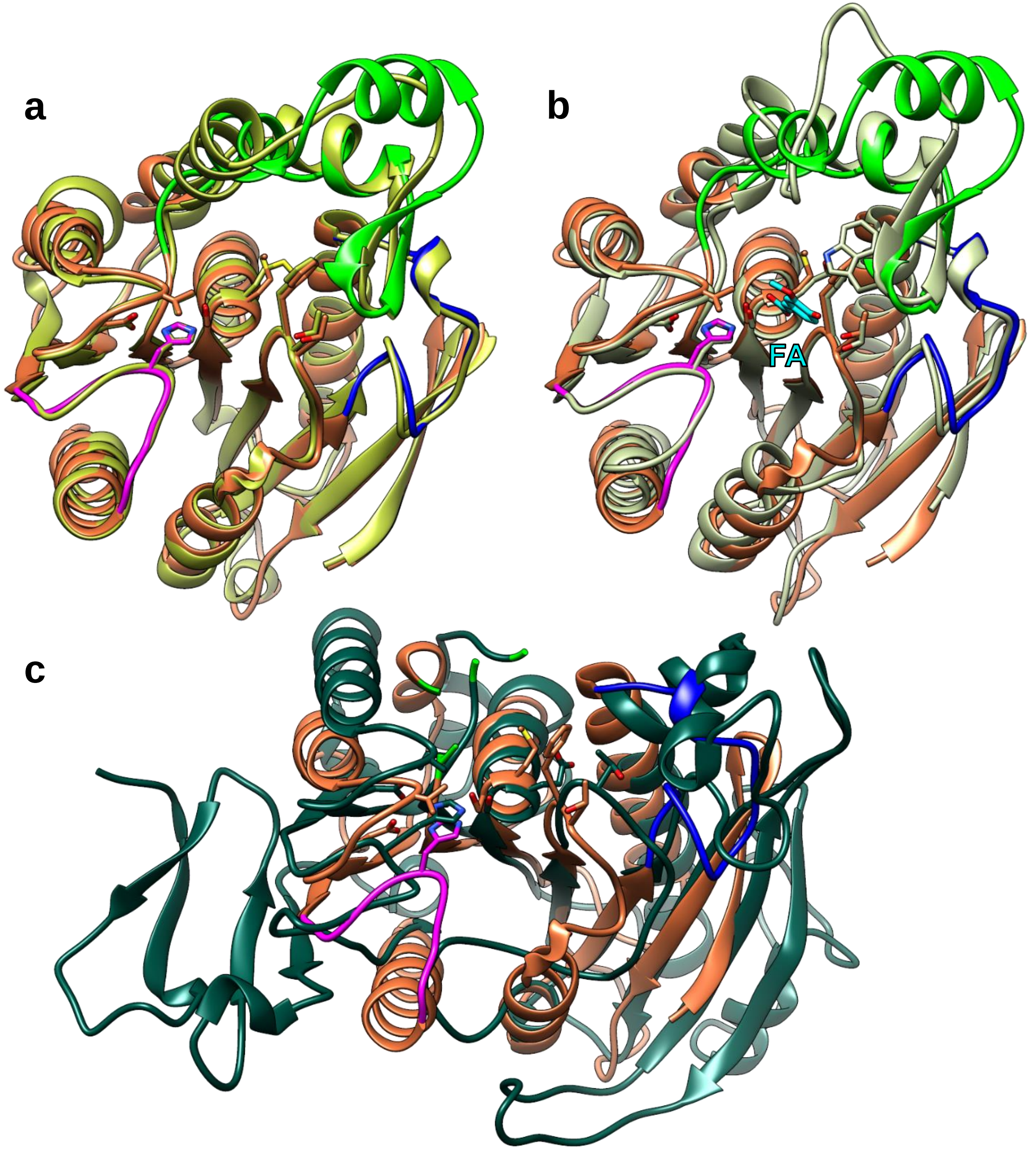


**Supplementary Fig. S4: The archaeal PETase PET46 is structurally homolog to ferulic acid esterases (FAEs).** The crystal structure of PET46 (coral orange) is compared to the crystal structures of GthFAE from *Geobacillus thermoglucosidasius* (lime green; PDB 7WWH; **a**), the Est1E FAE from *Butyrivibrio proteoclasticus* (cream white; PDB 2WTN) bound to ferulic acid (FA; cyan; **b**), and the tannase IsMHETase from *I. sakaiensis* (petrol green; PDB 6QZ4; **c**). The lid domains of PET46 and IsMHETase have been omitted in “c” for better visualization (bright green).

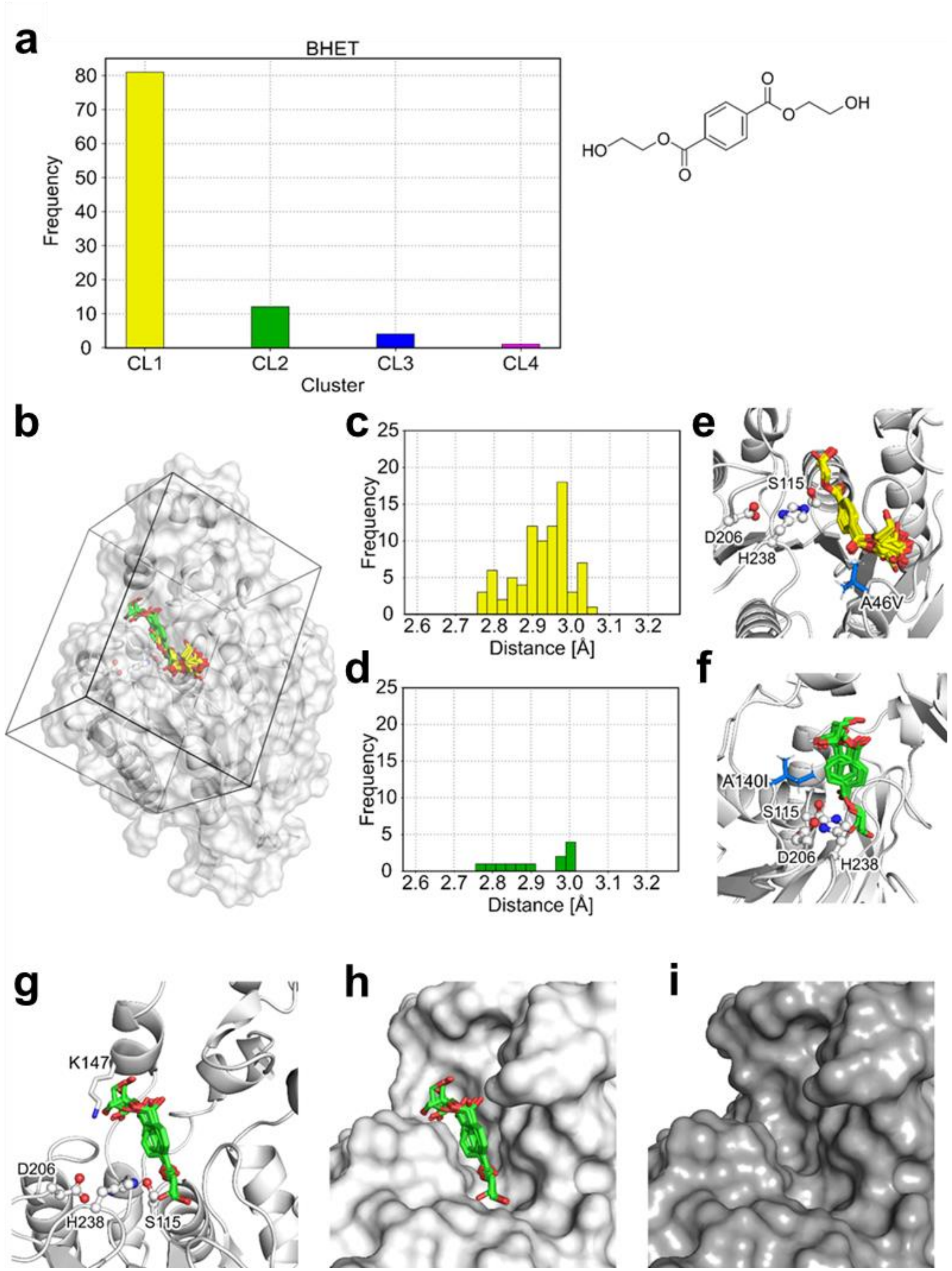


**Supplementary Fig. S5: Docking of BHET into PET46**. Docking of BHET yielded four possible binding poses (clusters CL1-CL4) in PET46 (**a**). Docked poses of the two largest clusters within PET46 with the box depicting the search space (**b**). Distributions of the smallest distances between the docked substrate's carbonyl carbon and the hydroxyl oxygen from the catalytic serine for the two largest clusters (**c**, **d**). Location of the substituted amino acids in the A46V variant (blue sticks, **e**), the A140I variant (blue sticks, **f**), and the K147 variant (white sticks, **g**) of PET46. The comparison of the substrate binding sites for the (H) WT (white surface, **h**) and the K147A variant (gray surface, **i**) shows an extended substrate binding site in the variant. The same orientation is used as for "g".

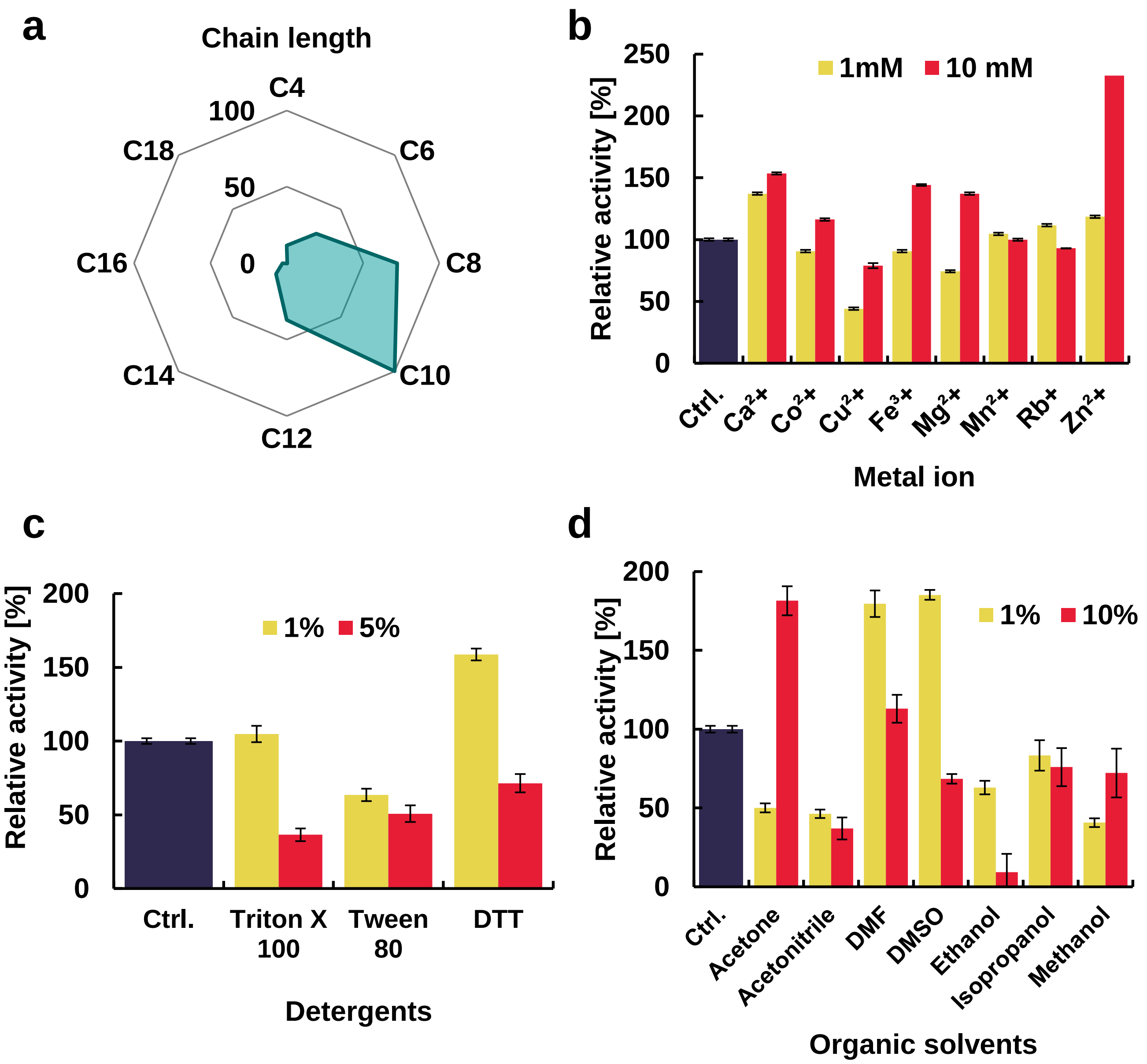


**Supplementary Fig. S6: Biochemical characterization of PET46.** Optimal *p*NP-ester acyl chain length was determined (**a**). The effect of metal ions (**b**), detergents (**c**) and organic solvents (**d**) on the activity of PET46 was studied compared to an additive-free control (Ctrl.). Error bars indicate the standard deviation of at least three replicates. Standard deviation in "a" was below 6 %.
